# Spatial and temporal expression of PORCN is highly dynamic in the developing mouse cochlea

**DOI:** 10.1101/2021.08.20.456516

**Authors:** Brianna L. Oliver, Caryl A. Young, Vidhya Munnamalai

**Author notes:** Correspondence should be addressed to: Vidhya Munnamalai, Ph.D., The Jackson Laboratory, 600 Main St., Bar Harbor, ME 04609, Ph. Equal contribution (Brianna L. Oliver, Caryl A. Young). Funding: NIH R21DC016376 and The Jackson Laboratory start-up funds (to VM). Declarations of interest: none.

## Abstract

The mammalian organ of Corti is a highly specialized sensory organ of the cochlea with a fine-grained pattern that is essential for auditory function. The sensory epithelium, the organ of Corti consists of a single row of inner hair cells and three rows of outer hair cells that are intercalated by support cells in a mosaic pattern. Previous studies show that the Wnt pathway regulates proliferation, promotes medial compartment formation in the cochlea, differentiation of the mechanosensory hair cells and axon guidance of Type II afferent neurons. WNT ligand expressions are highly dynamic throughout development but are insufficient to explain the roles of the Wnt pathway. We address a potential way for how WNTs specify the medial compartment by characterizing the expression of Porcupine (PORCN), an O-acyltransferase that is required for WNT secretion. We show PORCN expression across embryonic ages (E)12.5 - E14.5, E16.5, and postnatal day (P)1. Our results showed enriched PORCN in the medial domains during early stages of development, indicating that WNTs have a stronger influence on patterning of the medial compartment. PORCN was rapidly downregulated after E14.5, following the onset of sensory cell differentiation; residual expression remained in some hair cells and supporting cells. On E14.5 and E16.5, we also examined the spatial expression of *Gsk3β*, an inhibitor of canonical Wnt signaling to determine its potential role in radial patterning of the cochlea. *Gsk3β* was broadly expressed across the radial axis of the epithelium; therefore, unlikely to control WNT-mediated medial specification. In conclusion, the spatial expression of PORCN enriches WNT secretion from the medial domains of the cochlea to influence the specification of cell fates in the medial sensory domain.

**Highlights:**

- Wnt ligands are broadly expressed during cochlear development.
- PORCN expression is highly dynamic during early cochlear development
- PORCN becomes restricted to the medial domains along the longitudinal axis.
- Wnt medial specification is regulated at the level of WNT ligand secretion.

## 1. Introduction

The mammalian cochlea is a coiled, intricately patterned sensory organ informed by positional information during development. The organ of Corti (OC) houses mechanosensory hair cells (HCs), which convert mechanical energy from sound waves into auditory neural signals that are transmitted to the brain. There is a dichotomy within the OC with two domains: the medial sensory (MS) domain containing one row of inner hair cells (IHCs) that detect sound and the lateral sensory (LS) domain containing three rows of outer hair cells (OHCs) that amplify sound (Fig. 1). The inside of the coiled cochlea facing the modiolus, or the central axis of the cochlea, is referred to the medial side of the cochlea. The longitudinal axis communicates frequency selectivity, while the radial axis specifies neural processing by afferent and efferent projections. HC differentiation occurs in a base to apex direction beginning on approximately embryonic day (E)15 in the mouse cochlea (Kelley, 2006). Morphogen gradients endow cells in the cochlear epithelium with positional information to determine cell fate choices, giving rise to the complex organization of the cochlea (Das et al., 2012; Hayashi et al., 2008; Jacques et al., 2007; Mann et al., 2014; Munnamalai and Fekete, 2020; Munnamalai et al., 2012; Munnamalai et al., 2017; Ohyama et al., 2010; Urness et al., 2015). One major morphogen signaling pathway that establishes positional information in the cochlea is the Wnt pathway (Munnamalai and Fekete, 2013).

**Figure 1:**
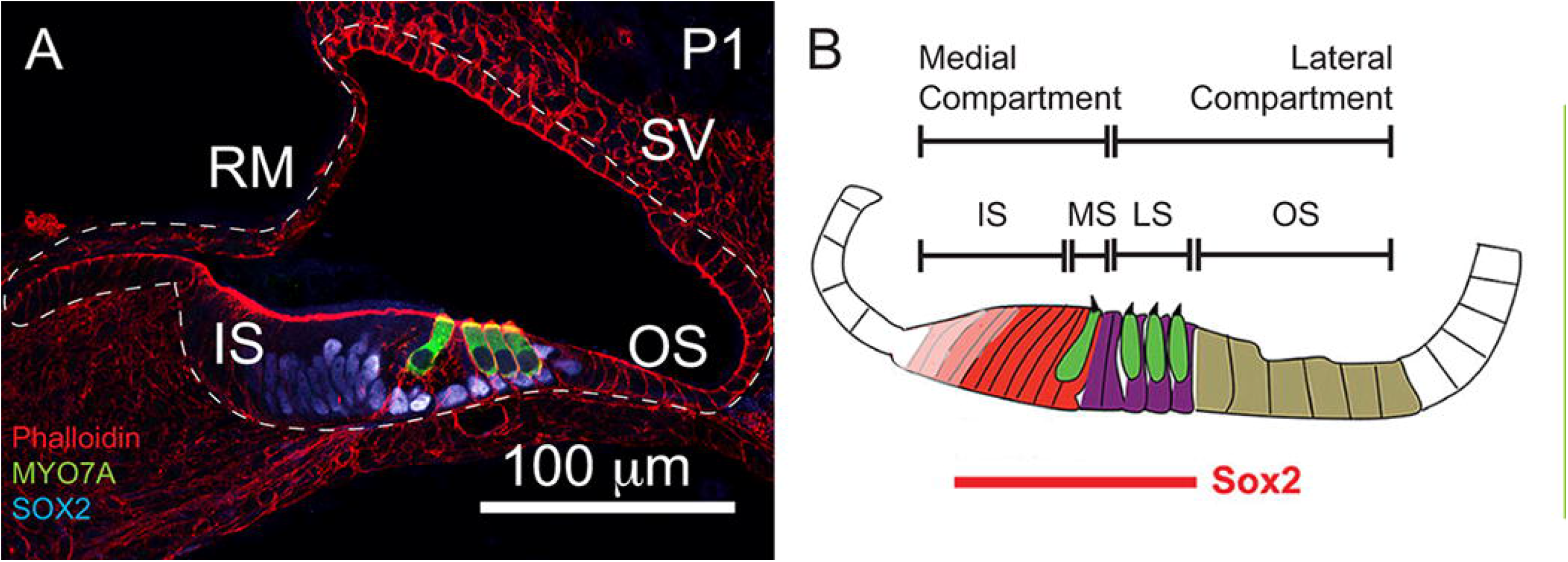
Organization of the Cochlea. (A) Phalloidin (red) labeling reveals the overall structure of the Scala media that houses the organ of Corti. SOX2 (light blue) labels the sensory domain and MYO7A labels the mechanosensory hair cells (green) in of the cochlea. (B) Cartoon depicting the domains on the floor of the cochlear duct. The inner hair cells and their support cells lie within the medial sensory (MS) domain and the outer hair cells and their support cells lie within the lateral sensory domain (LS). The organ of Corti is flanked by the inner sulcus (IS) and the outer sulcus (OS).

The Wnt pathway is known to regulate proliferation, differentiation, planar cell polarity, Ca^2+^ signaling and axon guidance of type II neurons in the cochlea (Dabdoub et al., 2003; Ghimire and Deans, 2019; Landin Malt et al., 2020; Macheda et al., 2012; Munnamalai and Fekete, 2013; Qian et al., 2007). In the canonical pathway secreted WNTs bind to frizzled receptors, inhibiting GSK3β. This allows β-Catenin to translocate into the nucleus and bind TCF/LEF transcription factors, activating gene transcription. Previous studies demonstrate canonical Wnt activity in the prosensory domain of the E12.5 cochlea (Jacques et al., 2012). At this stage, canonical Wnt signaling is important for regulating the proliferation of progenitors in the prosensory domain (Jacques et al., 2012; Munnamalai and Fekete, 2016) and radial patterning (Jansson et al., 2019; Shi et al., 2014). There are four *Wnts* expressed in the cochlea: *Wnt4, Wnt5a, Wnt7a*, and *Wnt7b* (Bohnenpoll et al., 2014; Munnamalai and Fekete, 2016; Qian et al., 2007). However, *Wnt5a, Wnt7a, and Wnt7b* are expressed on the floor of the duct, where the sensory domain is formed. Evidence suggests that Wnt activity directly regulates *Atoh1* expression, which is a master regulator for hair cell formation (Bermingham et al., 1999; Munnamalai and Fekete, 2016; Shi et al., 2010; Shi et al., 2014). However, previous studies show Wnt pathway activation promotes an expansion of the domain that houses IHCs and tall hair cells (THC) in the mouse and chicken cochleas respectively (Munnamalai and Fekete, 2016; Munnamalai et al., 2017). How this specificity is attained is unknown.

A recent study in the chicken otocyst showed that there is a dose dependent specification of sensory domain in the inner ear (Zak and Daudet, 2021). A similar dose-dependent specification of the medial compartment may be active in the developing cochlea. In order to identify a potential avenue for how the Wnt pathway specifies the medial compartment in the cochlea, we present a comparative study of the expression patterns of *Wnt ligands, Wntless (Wls)*, Porcupine (PORCN), and the Wnt inhibitor, GSK3β at different stages of cochlear development. We show that the spatial and temporal expression pattern of the PORCN enzyme alone, which palmitoylates WNT ligands, supports a mechanism for Wnt specification of medial fates in the developing cochlea by providing a spatial context for WNT secretion. The expression patterns of *Wls*, a G-protein coupled receptor that chaperons WNTs intracellularly for WNT secretion (Das et al., 2012), suggest that *Wls* is not a mechanism controlling medial domain specification. Thus, the cells in the domains from which WNTs are secreted, are exposed to a high dose of WNT ligands relative to the cells that are distal to PORCN-positive regions.

## 2. Results

In order to understand how the Wnt pathway specifies the medial domain, we sought to characterize and compare the expression of *Wnts* to the expression of *Wls* and PORCN in the cochlea. We examined the spatial expression by *in situ* hybridization of known *Wnt ligands, Wnt7b, Wnt7a*, and *Wnt5a*, which are present on the floor of the cochlear duct on E12.5, E14.5 and E16.5 (Fig. 2). On E12.5, *Wnt7b* was broadly expressed across the radial axis with a slight enrichment along the lateral edge of the cochlea, towards the outer sulcus (Fig. 2A). By E14.5, *Wnt7b* was homogenously expressed in the medial and lateral compartments along the longitudinal axis (Fig. 2Bi-Biii); however, by E16.5 *Wnt7b* was downregulated in the prospective sensory epithelium and this pattern was consistent along the longitudinal axis from base to apex (Fig. 2Ci-Civ). On E12.5, W*nt7a* was broadly expressed at very low levels (Fig. 2D). On E14.5, *Wnt7a* expression increased and was broadly expressed across the radial and longitudinal axes (Fig. 2Ei-Eiii). Similar to *Wnt7b*, by E16.5, *Wnt7a* was downregulated in the OHC region (Fig. 2Fi-Fiv). On E12.5, *Wnt5a* was expressed on the medial side and the lateral edge of the cochlea (Fig. 2G). By E14.5 *Wnt5a* expression was further defined in the medial sensory side and the lateral non-sensory side of the cochlea that precedes the *stria vascularis*. This expression pattern continued along the length of the cochlea (Fig. 2Hi-Hiii) (Qian et al., 2007). On E16.5, *Wnt5a* expression remained in the inner sulcus, but was down regulated in MYO7A-positive HCs (Fig. 2Iii, arrowheads) and continued along the cochlea (Fig. 2Ii-Iiv). Curiously, transcripts for all three *Wnt ligands* were spatially downregulated in the sensory domain and showed consistent expression along the longitudinal axis (Fig. 2C, F, I). *Wnt4* was expressed on the non-sensory side on E12.5 and from E14.5 onwards, *Wnt4* was restricted to the future Reissner’s membrane. The expression pattern was consistent along the longitudinal axis (data not shown). Based on this expression pattern, WNT4 is unlikely to play a direct role on medial specification.

**Figure 2:**
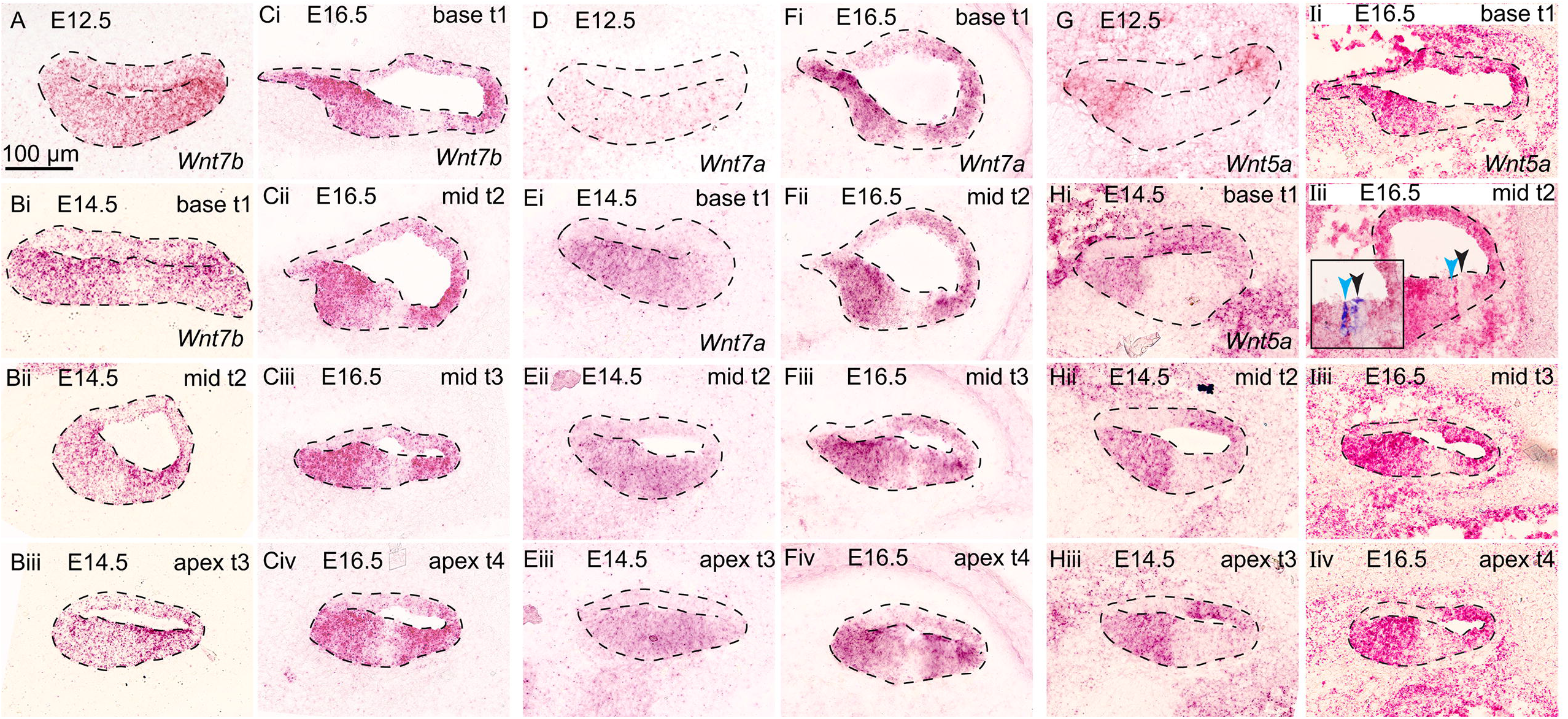
Cochlear expression of *Wnt* ligands on E12.5, E14.5 and E16.5. (A-C) *Wnt7b* was broadly expressed on E12.5 and E14.5 but was decreased in the prospective sensory epithelium on E16.5. (D-F) *Wnt7a* expression gradually increased from E12.5 to E16.5. On E16.5, *Wnt7a* was also downregulated in the sensory epoithelium (G-I) *Wnt5a* expression increased from E12.5 to E16.5 and was enriched in the medial domain and lateral non-sensory, or future stria vascularis. (I) Inset-MYO7A-positive HCs are labeled in dark blue. Light blue arrowhead labels in IHCs, and black arrowhead labels the OHCs. Cochleas in figures are oriented such that the medial side of the cochlea is placed on the left side and the lateral side of the cochlea is placed on the right. The expression patterns of all *Wnts* appeared to be similar along the longitudinal axis. Sample size of cochleas range from n = 4 to n = 6.

To understand the spatial regulation of WNT secretion in the cochlea, we examined the expression of *Wls* and *Porcn* by *in situ* hybridization on E12.5, E14.5 and E16.5 (Fig. 3). On E12.5 *Wls* was broadly expressed across the epithelium (Fig. 3A). On E14.5 *Wls* remained broadly expressed across the radial and longitudinal axes (Fig 3B). In the mid-turn and apex *Wls* was slightly enriched in the lateral half of the cochlea (Fig. 3Bii, Biii). By E16.5 *Wls* expression appeared to be comparatively lower, but homogeneous; however, it is spatially downregulated in the sensory domain (Fig. 3C). The *Wls* expression pattern was consistent along the longitudinal axis (Fig. 3Ci-Civ). A recent study characterized the expression of *Porcn*, by *in situ* hybridization, to show very low expression in the apex of the cochlea on E16.5 (Najarro et al., 2020). However, the probe used detected the *Porcn-a* isoform only, while there are four different isoforms for *Porcn* in total (Proffitt and Virshup, 2012). We used the same probe in conjunction with *Wls* probe and found that *Porcn-a* transcripts were barely detectable and significantly fewer compared to *Wls* transcripts in the epithelium on E12.5, E14.5 and E16.5, but was expressed at significant levels in the spiral ganglion neurons (SGN) (Fig. 3).

**Figure 3:**
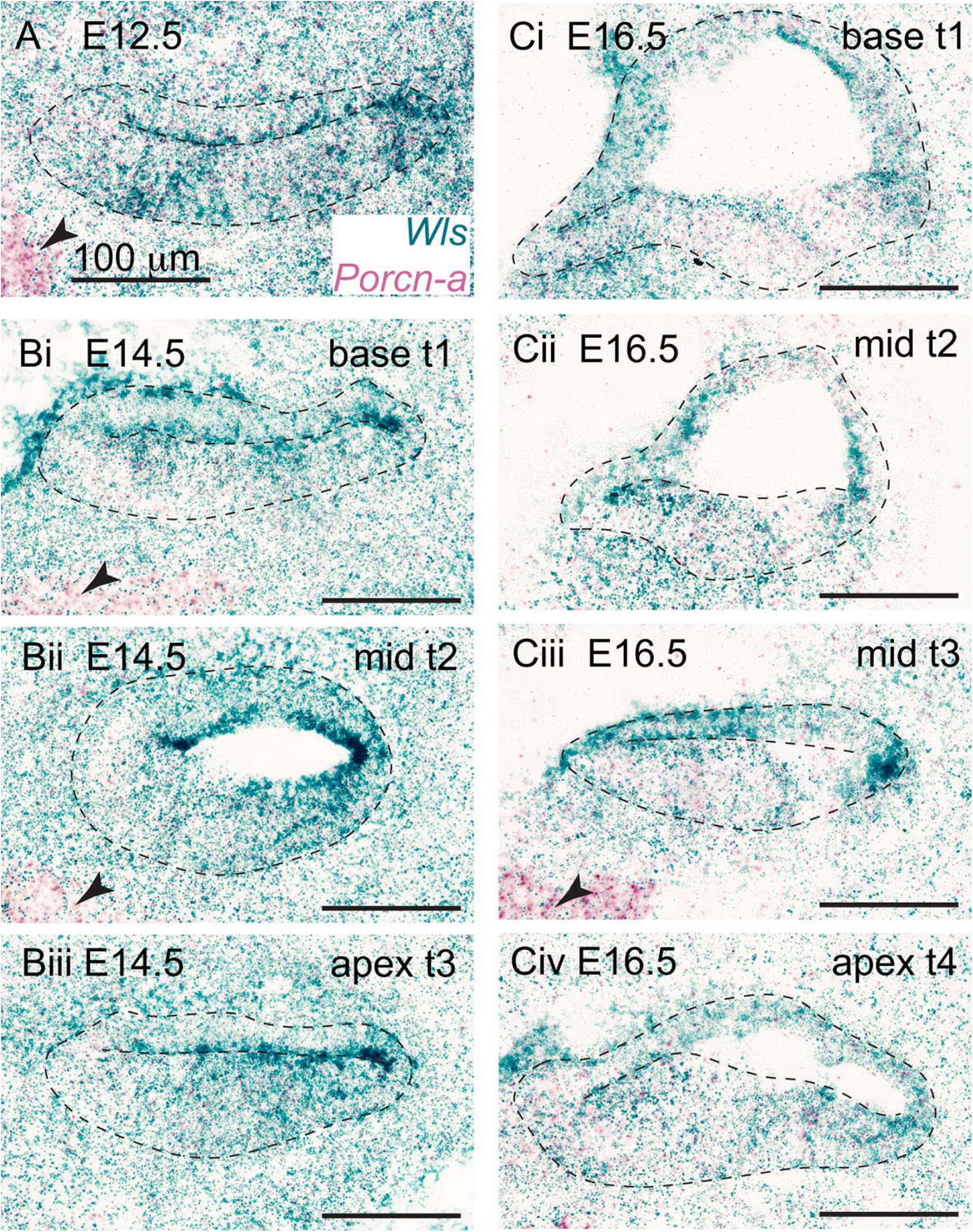
Cochlear expression of *Wls* and *Porcn-a* on E12.5, E14.5 and E16.5. (A) *Wls* was broadly expressed on E12.5, while *Porcn-a* was detected in low levels in the epithelium and moderate expression in the ganglion (black arrowhead). (B) On E14.5 *Wls* was broadly expressed but enriched laterally. *Porcn-a* was undetectable in cochlear epithelium but was observed in the ganglion (black arrowhead). (C) On E16.5 *Wls* was spatially downregulated in the sensory domain. *Porcn-a* was barely detectable within the epithelium but remained present in the ganglion (black arrowhead). Sample size of cochleas range from n = 3 to n = 4.

Given the spatiotemporal expression patterns of *Wnts* in the cochlea, we expected PORCN to be expressed in all corresponding domains to facilitate WNT secretion and that there must be other isoforms that are present. We used a synthetic peptide antibody that recognizes the sequence: ACRLLWRLGL PSYLKHASTV AGGFFSLYHF FQLHMVWVVL LSLLCYLVLF. A BlastP analysis for this sequence showed identity matches for the A, B and C isoforms. Our RNA sequencing performed on E14.5 cochleas did not detect the D isoform (data not shown). Therefore, we sought to determine the spatial expression of PORCN by immunolabeling. To test the specificity of the anti-PORCN antibody, we generated E14.5 *Isl1Cre; Porcn; Tdt* conditional knockout (*cKO)* embryos and PORCN was immunolabeled in both control and *Porcn cKO* cochleas (Fig. 4). TD-Tomato (TDT) expression reported the efficiency of Cre recombination under *Isl1Cre* expression. In control *Isl1Cre*^*+/-*^; *Porcn (X*^*flox*^ *X*^*+*^*); Tdt*^*+/-*^ cochleas, PORCN was medially enriched (Fig. 4A). In *Isl1Cre*^*+/-*^; *Porcn (X*^*flox*^ *Y); Tdt*^*+/-*^ *cKO* cochleas, PORCN immunolabeling was abolished (Fig. 4B). Consistent with the conditional deletion of *Porcn*, TDT expression showed that Cre recombination occurred across the floor of the duct (Fig. 4C, D). This validated the specificity of the anti-PORCN antibody. *Jag1* is a predicted Wnt target gene and has shown to be a direct Wnt target in hair follicles (Estrach et al., 2006). Although, this has not been demonstrated in the inner ear and *Jag1* responsivity to Wnt signaling changes throughout development in the otocyst and the cochlea (Munnamalai and Fekete, 2016; Zak and Daudet, 2021), we analyzed the JAG1 domain in *Porcn cKOs*. In control cochleas, JAG1 was expressed medially within the PORCN domain (Fig. 4A, E). The global loss of WNT ligands in *Porcn cKO* cochleas completely abolished JAG1 (Fig. 4F). This validated both the anti-PORCN antibody and the *Porcn cKO* mouse model.

**Figure 4:**
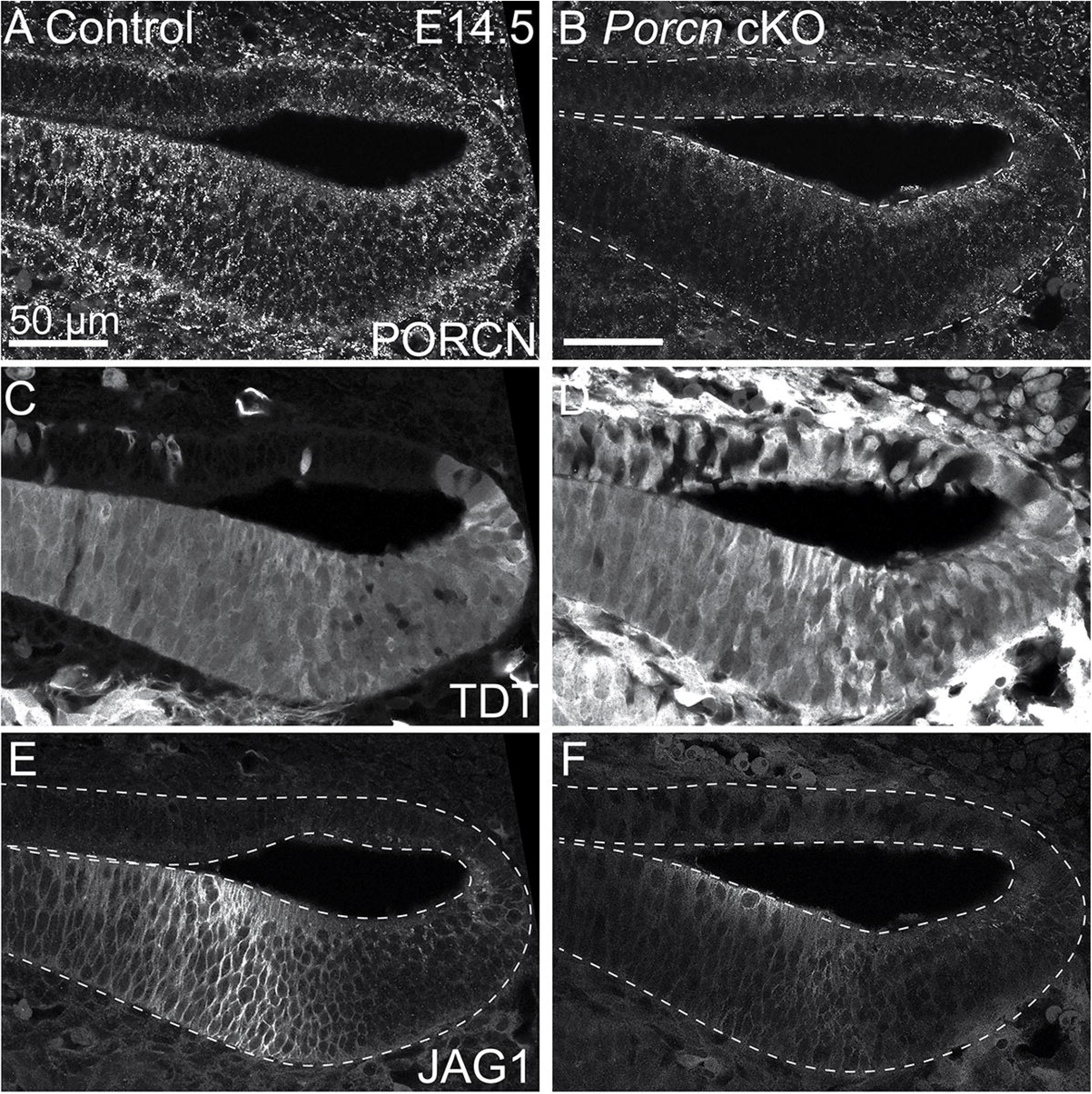
Validation of the anti-PORCN antibody on E14.5 in *Isl1Cre; Porcn cKO* cochlea. (A) PORCN was enriched in the medial domain. (B) PORCN expression was abolished in *Porcn* cKO cochlea. (C-D) TDT expression reported efficient Cre recombination on the floor of the duct. (E) Expression of the Wnt target JAG1 overlapped with PORCN expression in the medial domain. (F) JAG1 expression was abolished in *Porcn* cKO cochlea. n = 8 cochleas.

We then characterized the spatio-temporal expression of PORCN relative to SOX2, which labels the prosensory domain or support cells, on E12.5-E14.5 (Fig. 5), and on E16.5 and postnatal day (P)1 (Fig. 6). On E12.5, PORCN was broadly expressed across the radial axis and enriched centrally in the floor of the duct, adjacent to the medial SOX2 domain (Fig. 5A). On E13.5, PORCN expression sweeps from base to apex along the longitudinal axis. SOX2 expression was highest in the base similar to E12.5 and reduced towards the apex (Fig. 5B). In the mid-turn (t2), the SOX2-positive prosensory domain was shifted centrally with low levels of PORCN on the medial edge. In the apical turn (t3), PORCN was not yet expressed (Fig. 5B, B’). On E14.5, PORCN expression remained broad in the basal turn (t1) (Fig. 5C); however, in the mid-turn, t2 PORCN was significantly confined to the medial domain adjacent to SOX2 (Fig. 5D). PORCN expression was lowest in t3 but enriched in the medial region adjacent to the SOX2 domain (Fig. 5E). Thus, PORCN expression was consistently broad in the basal turns from E12.5-E14.5. However, further along the longitudinal axis, PORCN became restricted medially adjacent to the sensory domain. Although there was a base to apex sweep, the spatial expression of PORCN was always restricted to the medial side in the mid and apical turns of the cochlea.

**Figure 5:**
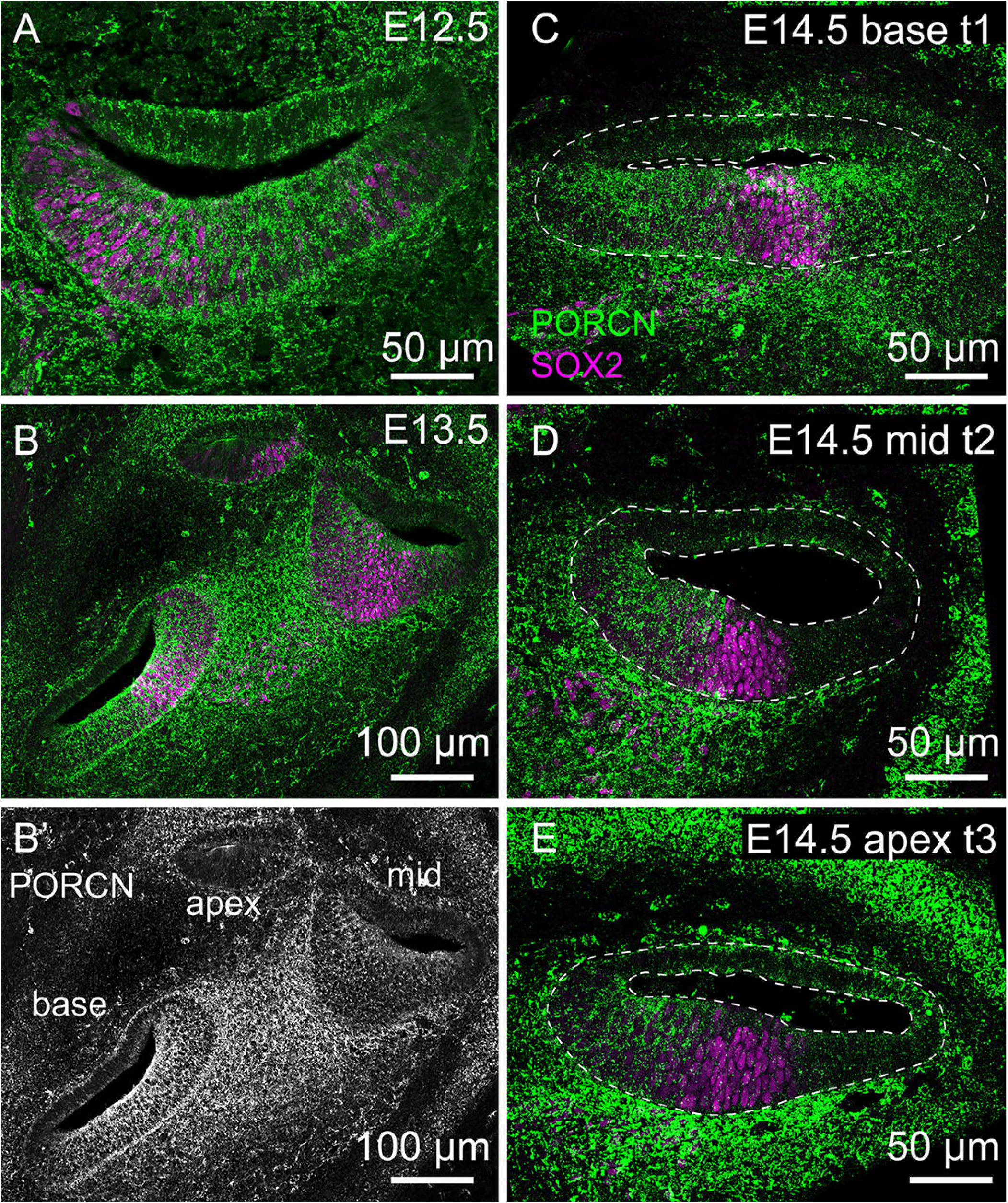
PORCN expression on E12.5, E13.5 and E14.5 relative to the SOX2 sensory domain. (A) On E12.5, PORCN expression (green) was broad in the basal turn of the cochlea. (B, B’) On E13.5, PORCN swept in a basal-to-apical gradient. (C-E) On E14.5, PORCN expression was highest in the base across the radial axis, but then became restricted to the medial domain towards the apex. (A-E) SOX2 expression (magenta). Sample size of cochleas range from n = 3 to n = 10.

**Figure 6:**
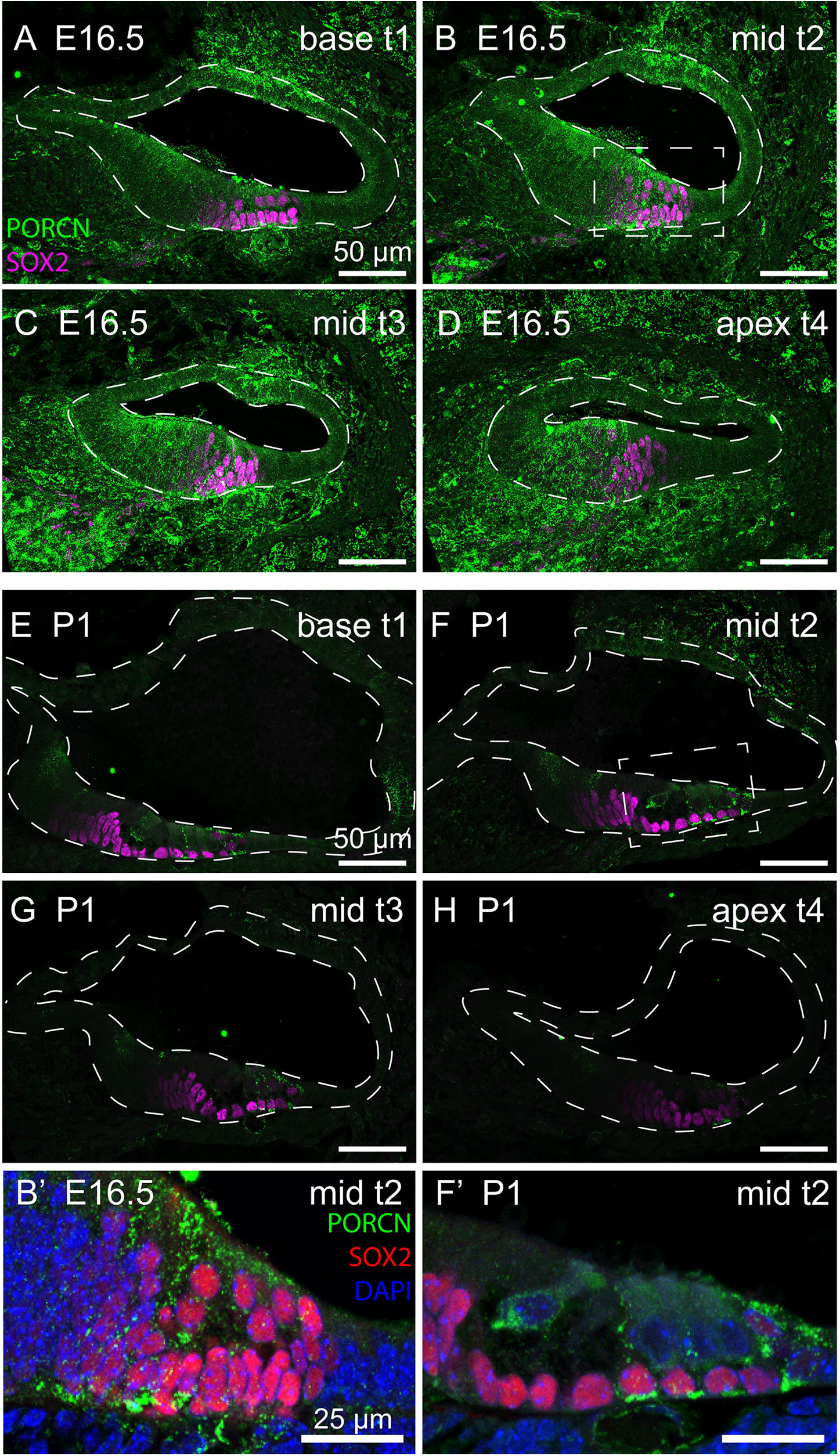
PORCN expression on E16.5 and P1 relative to the SOX2 sensory domain. (A-D) On E16.5, PORCN (green) expression remained in the sensory domain in the base but was restricted to the medial domain towards the apex. (B’) Inset shows that PORCN was enriched in the domain containing the IHCs. (E-H) On P1, PORCN expression was drastically decreased in the cochlea. (F’) Inset shows PORCN expression was retained in the IHCs, Deiters’ cells and Hensen’s cells. (A-H) SOX2 expression (magenta). (B’, F’) SOX2 expression (red), DAPI (blue). Sample size of cochleas range from n = 3 to n = 4.

Compared to E14.5, PORCN was dramatically reduced on the medial side by E16.5 (Fig. 6). In t1 and t2 there was very low levels of PORCN in the IHC and its neighboring supporting cell (Fig. 6A, B, B’). Unlike E14.5, there was no PORCN expression in the far medial domain adjacent to SOX2 (Fig. 6A, B). In t3, enriched PORCN expression remained in the medial compartment, but there was low level PORCN expression within the SOX2 domain (Fig. 6C). On E16.5, the PORCN expression pattern in the apical turn, t4, was similar to the E14.5 mid-turn, t2 with an enrichment in the medial domain adjacent to the SOX2 domain (Fig. 5D, 6D). Towards the apex, PORCN expression never extended across the radial axis (Fig. 5D, 5E, 6D), unlike the basal turns from E12.5-E14.5 (Fig. 5A, B, C). By P1, PORCN was drastically reduced along the entire longitudinal axis (Fig. 6E-H). In t1 and t2, PORCN was observed on the basolateral side of the IHCs, and in the Deiters’ and Hensens’ cells (Fig. 6E, F, F’). Towards the apex, PORCN was barely detectable (Fig. 6G, H).

In previous studies, pharmacological activation of the Wnt pathway using a GSK3β inhibitor, CHIR99021 (CHIR), promoted the formation of the medial sensory domain that houses the IHCs on approximately E13.5-E14.5 (Munnamalai and Fekete, 2016). To address whether GSK3β, which is further downstream of PORCN, influences Wnt-mediated medial specification on E14.5, we examined the spatial expression of GSK3β across the radial axis. *In situ* hybridization for *Gsk3β* transcripts on E14.5 showed that *Gsk3β* was ubiquitously expressed across the epithelium (Fig. 7A). On E16.5, *Gsk3*β was slightly downregulated in the region medial to the IHCs, but still expressed at significant levels (Fig. 7B). Immunolabeling of E14.5 cochleas for GSK3β, supports our *in-situ* hybridizations showing homogenous expression across the radial axis at the same age (Fig. 7A, C). To validate the specificity of the anti-GSK3β antibody, we immunolabeled for GSK3β in E14.5 *Isl1Cre; Gsk3βcKOs* cochleas (Fig. 7D). On E14.5, GSK3β expression was completely absent in *Gsk3*β *cKOs* cochleas and the spiral ganglion (SGN) and is accompanied by a slightly expanded SOX2 domain compared to controls (Fig 7C, C’ D, D’) as previously described (Munnamalai and Fekete, 2016). This validated the specificity of the anti-GSK3β antibody and confirmed that on E14.5, GSK3β is homogenously expressed across the radial axis.

**Figure 7:**
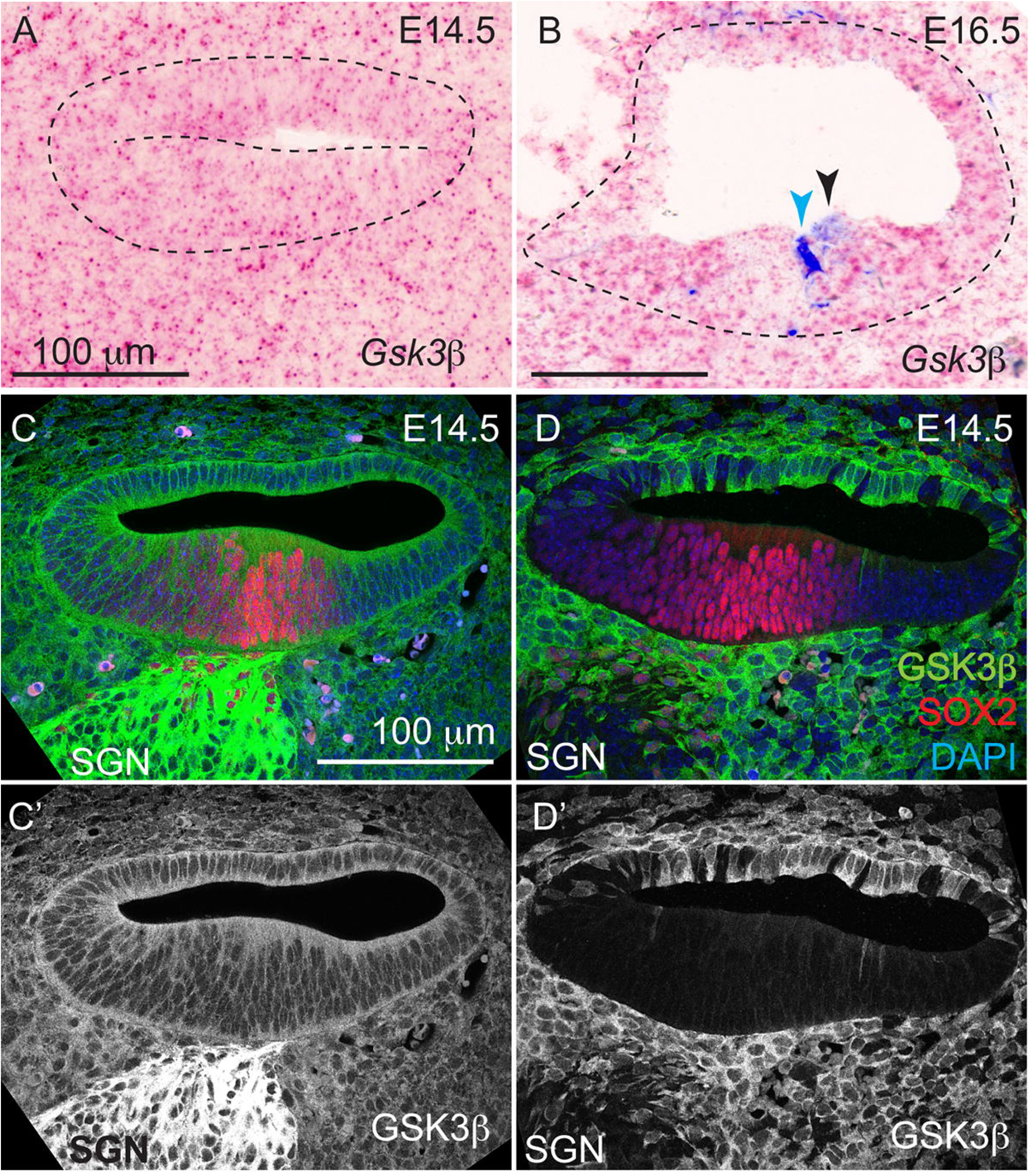
GSK3β cochlear expression on E14.5 and E16.5. (A) *In situ* hybridization for *Gsk3β* on E14.5 showed ubiquitous expression in the cochlea. (B) On E16.5, *Gsk3β* was enriched on the medial sensory and lateral non-sensory domains. MYO7A-positive HCs are labeled in dark blue. Light blue arrowhead labels in IHCs, and black arrowhead labels the OHCs. (C) On E14.5, GSK3β was ubiquitously expressed in the cochlea similar to *Gsk3β* transcript expression on E14.5, the surrounding mesenchyme and the spiral ganglion (SGN). (D) Anti-GSK3β immunolabeling was absent in *Isl1Cre; Gsk3β cKO* cochlea and the SGN, validating the anti-GSK3β antibody. (C’-D’) Immunolabeling of the SOX2 sensory domain in control and *Gsk3β* cKO cochlea for comparison. Sample size of cochleas range from n = 4 to n = 6.

## 3. Discussion

In the developing cochlea, the Wnt pathway plays dual roles, i.e., the regulation of cell cycle proliferation and differentiation (Dabdoub et al., 2003). Three *Wnt ligands* are known to be expressed at significant levels on the floor of the duct to influence the formation of the sensory domain: *Wnt5a* (Qian et al., 2007), *Wnt7a* (Dabdoub et al., 2003) and *Wnt7b* (Bohnenpoll et al., 2014; Munnamalai and Fekete, 2016). In both chicken and mouse cochleas, over-activation of Wnt signaling at the ligand level and by GSK3β inhibition showed that Wnt activation promotes the medial, or neural compartment that gives rise to IHCs (Munnamalai and Fekete, 2016; Munnamalai et al., 2017). This specificity is surprising given the cumulative expression patterns of WNT ligands. During the early developmental stages between E12.5-E14.5, there is a Wnt ligand that is transcribed within any given region of the cochlea. What determines the specificity of this medial identity in the cochlea?

Immunolabeling for PORCN during the early developmental stages between E12.5 and E14.5 showed a broad but consistent expression towards the basal end of the cochlea that was restricted to the medial domains towards the apex (Fig. 5). On E16.5 the PORCN expression seems meaningless for WNT secretion as there appears to be no *Wnt-ligand* expression within HCs. Future studies would benefit from identifying the spatial expression of all the PORCN isoforms that are expressed in the cochlea.

The differences in expression patterns along the longitudinal axis suggests a possible role for regulating tonotopy, leaving open the pursuit of an intriguing future direction. This longitudinal expression pattern may also be responsible for influencing the pathfinding of type II neurons, which were affected in *Porcn cKOs* (Ghimire and Deans, 2019). Along the radial axis, particularly in the mid and apical turns, we saw an enrichment of PORCN in the medial domain adjacent to the SOX2-positive sensory domain (Fig 5. D, E). This observation supports a stronger role for Wnt signaling in promoting medial fates over lateral fates, particularly up to E14.5 while the medial domain is specified. There was a drastic downregulation of PORCN in the medial domain of the cochlea after E15/ E16.5, particularly on the base-end of the cochlea (Fig. 6A, E-H). Thus, Wnt signaling is significantly downregulated after E15 in a normal context. Towards the apex, however, PORCN expression was reminiscent of earlier stages of development on E14.5 (Fig. 5D, 6C, 6D). This suggests that once sensory cells have initiated differentiation, WNT secretion is downregulated. It is curious as to why *Wnt* transcripts are so abundantly expressed, when its major secretory component is decreased. Interestingly, PORCN expression was retained in the IHCs, Deiters’ cells and Hensen’s cells (Fig. 6B’, F’). Although Wnt activation increases *Fgf8*-positive IHCs (Munnamalai and Fekete, 2016), and THC fates (Munnamalai et al., 2017), loss-of-function Wnt mutant studies do not clearly address whether Wnt signaling is required for HC-type specification. One explanation could be the inability to finely titrate the time window for abolishing Wnt function in transgenic animals. *Wls* is also important for WNT secretion. Although there was a slight enrichment on lateral half of the cochlea on E14.5, there are abundant *Wls* transcripts expressed on the medial half to facilitate WNT secretion; thus, PORCN would still be the limiting factor for regulating secretion.

To address whether GSK3β, which is downstream of WNT ligands, influences radial identity in either a Wnt-dependent or independent manner, we characterized its spatial expression on E14.5 and E16.5. On E14.5, the homogenous expressions of GSK3β protein and *Gsk3β* transcripts across the radial axis suggest that there is no radial influence for GSK3β signaling on E14.5 and that Wnt-mediated medial compartment specification in the cochlea is indeed regulated at the ligand level by PORCN by E14.5. On E16.5, there was a slight decrease of *Gsk3β* in the region medial to the IHCs (Fig. 7B). However, by E16.5, the IHCs have already been specified, so WNT secretion to specify medial compartment cell fates isn’t required. *Gsk3β* was expressed in the extreme medial and lateral edges, coincident with a drastic downregulation of PORCN in the medial compartment, which further supports that canonical Wnt signaling is ‘turned off’ in the medial domains. However, there is a potential for PORCN secretion; hence Wnt activity to be present in the sensory domain and for GSK3β to influence Wnt-independent kinase signaling in a radial manner on E16.5. Wnt inhibitors such as SFRPs, WIF1 and DKKs are also expressed in the cochlea to influence cell fate decisions across the radial axis (Geng et al., 2016). *Sfrp1* expression remained broad from E12.5 to P0, after which it was downregulated in the sensory domain. Previous studies show that *Sfrp3*, also known as *Frzb*, was consistently expressed in the OS (Geng et al., 2016; Qian et al., 2007), but postnatally it was expressed in the OHCs. *Sfrp2* in contrast, was consistently expressed at very low levels from E12 to P0 (Geng et al., 2016). *Kremen1* was expressed in the sensory domain on E15.5 and in later ages (Mulvaney et al., 2016). *Dkk3* was expressed in the GER medial to the IHCs after E15.5. With the exception of *Dkk3* after E15.0, most inhibitors appear to be expressed on the lateral half, which further influences the specification of medial cell fates. DKK3 and KREMEN1 likely have an impact on E15.5, after the radial domains have been specified between E12.5-E14.5. In conclusion, the spatial expression of PORCN provides some level of control for Wnt pathway specification of the medial domains in the developing mammalian cochlea during early embryonic development on E12.5-E14.5.

## 4. Methods

### Mice

*B6*.*119(Cg)-Gsk3β* ^*tm2Jrm*^*/J* (*Gsk3β* ^*flox*^) (Patel et al., 2008) mice (The Jackson Laboratory, JAX, Bar Harbor, Maine, USA) and *Porcn*^*flox*^ (Barrott et al., 2011) mice (Charles Murtaugh, University of Utah) were used to generate cKOs. *Porcn*^*flox*^ mice were crossed with *B6*.*Cg-Gt(ROSA)26Sor* ^*tm14(CAG-tdTomato)Hze*^*/J* (Madisen et al., 2010) (JAX) reporter mice to generate double-floxed *Porcn*^*flox*^; *Tdt* ^*flox*^ mice. *Porcn*^*flox*^ mice, or *Gsk3β* ^*flox*^ were crossed with *Isl1*^*tm1(Cre)Sev*^*/J* (*Isl1Cre*) mice (Yang et al., 2006) (JAX). The first day a plug was observed, was designated as E0.5. E14.5 embryos were harvested and fixed in 4% PFA for processing. Swiss Webster (SW) dams (Envigo, Indiana, USA) were time-mated and embryos, or pups were euthanized on E12.5, E14.5, E16.5 and P1. All animal procedures were performed in accordance with the Institutional Animal Care and Use Committee (IACUC) guidelines at The Jackson Laboratory.

### Histology

Embryos were decapitated and fixed overnight with 4% paraformaldehyde (PFA) (Electron Microscopy Sciences, Hatfield Pennsylvania, USA), and immersed in a series of 10%, 20%, and 30% sucrose solutions at 4 °C. Heads were embedded with Tissue Freezing Medium (TFM) (General Data Healthcare, Cincinnati, Ohio, USA) and cryo-frozen in liquid N_2_. Tissues were stored at −80°C and cryo-sectioned for immunofluorescence and *in situ* hybridization.

### Immunofluorescence

Cryo-sectioned tissues were blocked with 2% Donkey serum (Jackson ImmunoResearch, West Grove, Pennsylvania, USA), followed by overnight primary antibody incubation at 4□. Primary antibodies used: rabbit *α*-PORCN (Invitrogen, Waltham, Massachusetts, USA, PA5-43423), goat *α*-SOX2 (R&D Systems, Minneapolis, Minnesota, USA, AF2018), rabbit *α*-GSK3β (Cell Signaling, Danvers, Massachusetts, USA, 12456S). *α*-GSK3β antibody was previously used in a GSK3β study (Ellis et al., 2019). GSK3β antibody requires an antigen retrieval step. Tissues were incubated in a 10 mM sodium citrate buffer with 0.05 % Tween 20 (pH of 6) (Fisher Scientific) for 15 minutes at 99□ prior to fixation. Secondary antibody incubation was performed with Alexa-conjugated antibodies (Life Technologies Corporation, Eugene, Oregon, USA). Tissue was mounted with Fluoromount G mounting medium (Life Technologies Corporation).

### *In situ* hybridization

Cryo-sectioned tissue was fixed with 4% PFA at room temperature for one hour followed by H_2_O_2_ treatment and antigen retrieval (Advanced Cell Diagnostics, Newark, California, USA) at 99□ for 5 minutes. Probes used: *Wnt5a* (ACD, 316791), *Wnt7a* (ACD, 401121), *Wnt7b* (ACD, 401131), *Wls* (ACD, 405011), *Porcn-a* (ACD, 404971-C2) and *Gsk3β* (ACD, 458821). The protocol was followed according to the manufacturer’s instructions. When co-immunolabeling for HCs, after in situ hybridization, anti-MYO7A antibody, followed by secondary antibody incubations were performed following a regular immunofluorescence protocol.

### Microscopy

Images were acquired at 20X and 40X magnifications on a Zeiss LSM800 confocal microscope at The Jackson Laboratory Microscopy Core and a brightfield/ epifluorescence Olympus BX51 microscope with a Spot insight CMOS camera in the Munnamalai lab. Image stacks were analyzed on FIJI and images were processed in Adobe Photoshop and Illustrator.

## Acknowledgements

The authors gratefully acknowledge the contribution of the Microscopy Core at The Jackson Laboratory to generate the work described in this publication. The authors also thank Dr. Holly Beaulac in the Munnamalai lab for critical reading of this manuscript. This work was supported by the National Institutes of Health (R21DC016376) and start-up funds from The Jackson Laboratory to VM.

